# Suppressing Transfer of Antibiotic Resistance by a Small RNA Virus

**DOI:** 10.64898/2026.03.25.714153

**Authors:** Zachary Lill, Jirapat Thongchol, David Solis, Junjie Zhang

## Abstract

The global rise of antimicrobial resistance (AMR) demands innovative strategies to limit the spread of multidrug-resistant bacteria. Conjugative plasmids, particularly those in the incompatibility group P (IncP), play a central role in disseminating resistance genes across diverse bacterial species via their encoded Type IV secretion systems (T4SS). Here, we characterize the single-stranded RNA bacteriophage (ssRNA phage) PRR1, which selectively targets AMR ESKAPEE pathogens carrying the IncP plasmid RP4, and assess its ability to inhibit conjugation. Using cryo–electron microscopy, we first resolved the mature PRR1 virion at 3.45 Å resolution revealing two phage maturation protein (Mat)-RNA interactions within the 3’ untranslated region (UTR) – a conserved interaction (Mat-U1) and a novel interaction (Mat-V1) for ssRNA phages. To characterize the PRR1-RP4 pilus interaction, we performed alanine-scanning mutagenesis and pinpointed four critical TrbC pilin residues (S12, W13, S72, and R77) for infection. Computational modeling revealed that these residues are located near the termini of the pilin at the phage-pilus interface. Notably, native and non-infectious, UV-crosslinked PRR1 were sufficient to block RP4 transfer, indicating conjugation inhibition does not require a complete infection cycle. Finally, combining PRR1 and antibiotic treatment yielded nine unique phage-resistant mutants within T4SS-associated genes on the RP4 plasmid. Eight of these mutants nearly abolished conjugation, while the *trbE* frameshift mutant retained ∼30% of wild-type efficiency, which is pivotal to clarifying the relationship between phage infection and pilus function. Collectively, these results establish ssRNA phages as specific T4SS plasmid targeting agents and underscore their potential to limit horizontal gene transfer in AMR pathogens.

**IMPORTANCE:** Antimicrobial resistance (AMR) spreads rapidly through horizontal gene transfer, largely driven by conjugative plasmids. Despite their central role, few strategies exist to directly block plasmid transfer. Here, we show that the IncP plasmid-dependent ssRNA phage PRR1 can inhibit the spread of antibiotic resistance genes by targeting the RP4 T4SS pilus. Structural and mutational analyses reveal previously unrecognized RNA packaging interactions and identify four pilin residues critical for infection. Remarkably, non-infectious PRR1 particles alone are sufficient to block conjugation, offering inhibition without the selective pressure from phage replication. Almost all PRR1-resistant RP4 mutants lost or had severely reduced plasmid transfer, while the remaining mutant is critical for studying the link between T4SS function and phage infection. These results highlight ssRNA phages as precise agents for limiting AMR gene dissemination.

## INTRODUCTION

Antimicrobial-resistant (AMR) bacteria represent a growing global health crisis, with projections estimating over $1 trillion in annual healthcare costs and approximately 2 million deaths per year by 2050.^1,2^ While the threat is urgent, antibiotic innovation has stalled, and microbial resistance now outpaces the development of new drug classes. As a result, alternative strategies to combat AMR are urgently needed.

A group of seven bacterial pathogens, collectively known as the ESKAPEE (*Enterococcus faecium, Staphylococcus aureus, Klebsiella pneumoniae, Acinetobacter baumannii, Pseudomonas aeruginosa, Enterobacter* species, and *Escherichia coli*) pathogens, are primary drivers of nosocomial infections and are frequently associated with multidrug resistance (MDR).^3^ AMR can arise through spontaneous mutations or, more commonly, via horizontal gene transfer (HGT).^4^ Among HGT mechanisms, conjugative plasmids are particularly concerning, as they enable rapid interspecies transfer of resistance genes through Type IV secretion systems (T4SS). Notably, several bacteriophages (phages), including alphatectiviruses, inoviruses, and some fiersviruses, target T4SS, encoded on conjugative plasmids, as their receptors. These so-called plasmid-dependent phages (PDPs) often share the host range of the plasmids they exploit.^5^

Single-stranded RNA (ssRNA) phages in the Fiersviridae family, including MS2 and Qβ, are classical PDPs, which use the conjugative pilus encoded by the F-plasmid (F-pilus) as a receptor and deliver their genomes upon pilus retraction. Recent studies demonstrate that MS2 and Qβ can detach the F-pilus upon entry.^6^ Importantly, to resist PDPs’ infection, bacterial hosts usually lose the plasmids encoding T4SS, leading to depilated cells to which phage cannot absorb.^7^ This results in bacteria that can no longer spread antibiotic resistance genes due to the loss of T4SS. Therefore, PDPs can be potential tools for limiting the spread of antibiotic resistance.

Compared to the narrow-host-range F-plasmid system, primarily in enteric bacteria such as *E. coli*, the incompatibility group P (IncP) plasmid RP4 is known as one of the broadest-host-range conjugative plasmids, with transfer to Gram-negative, Gram-positive bacteria, and yeast observed.^8–10^ Critical to the growing crisis of AMR, it is commonly found in ESKAPEE pathogens and confers resistance to multiple antibiotics. The RP4 plasmid of the IncP family encodes the complete IncP T4SS, with all transfer (Tra) genes located in the Tra2 operon except *traF* in the Tra1 operon (**Fig. 1A-C**).^11–13^ There are three antibiotic-resistance genes on RP4 that resist tetracycline (*tet*), ampicillin (*bla*), and kanamycin (*aphA*), respectively. The ssRNA phage PRR1 specifically uses the RP4-encoded pilin protein, TrbC, as its receptor.^14–16^ This makes PRR1 unique compared to MS2 or Qβ, as PRR1 can interact with the RP4-encoded receptor to infect a wide variety of pathogenically relevant species, such as *P. aeruginosa, E. coli, S. typhimurium*, and *V. cholerae*, with the former two being members of the ESKAPEE pathogens.

**Figure 1.**
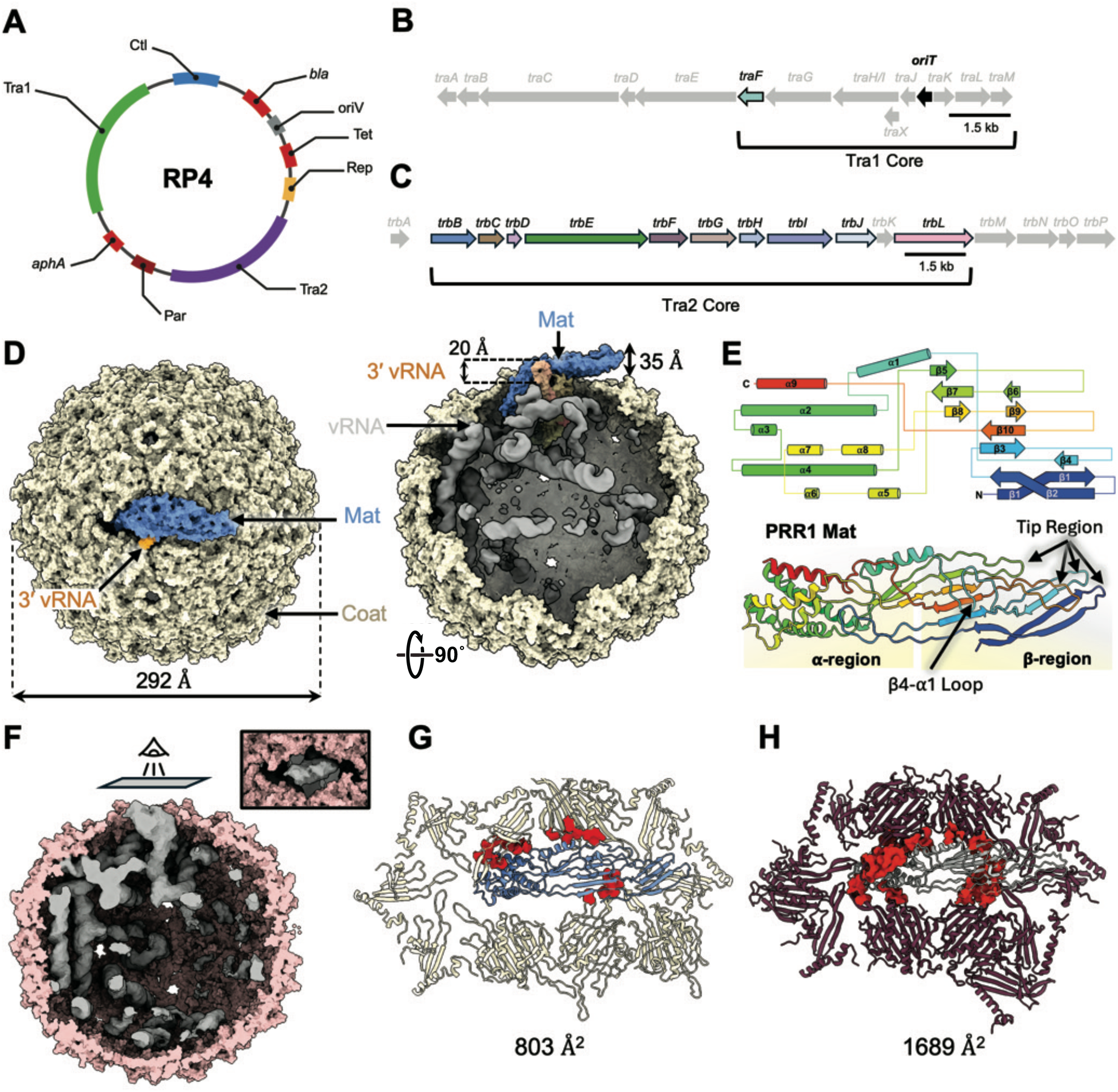
Organization of the RP4 plasmid and structural analysis of the PRR1 virion. **(A)** Organization of the major operons in the RP4 plasmid. Tra1 and Tra2 contain the genes for the relaxosome and T4SS, respectively. The Ctl operon helps regulate gene expression and the Rep operon functions in plasmid replication through initiation at the oriV site. The Par operon encodes the toxin-antitoxin host-defense system ParDE. Lastly, the three antibiotic resistance genes are encoded on *bla* (β-lactamase enzyme that breaks down ampicillin), *aphA* (aminoglycoside-3-phosphotransferase enzyme inactivating kanamycin), and the Tet operon (encodes *tetA* which produces the tetracycline efflux pump). **(B)** The genes on the Tra1 (transfer 1) operon are shown. Genes in gray are not critical for formation of the T4SS machinery and pilus biogenesis. The origin of transfer (oriT) is highlighted in black. **(C)** The genes on the Tra2 (transfer 2) operon are shown. Genes in gray are not critical for formation of the T4SS machinery or pilus biogenesis. **(D)** Cryo-EM reconstruction of the mature PRR1 virion, showing the Coat (tan), Mat (blue), and the viral RNA (vRNA, gray) with the 3′ end of the vRNA labeled (orange). The virion diameter (292Å) and Mat prominence (35Å) are labeled. One stem of the 3’ vRNA extends 20Å outside the capsid. Left: top-down view of the intact virion from the Mat. Right: cross-sectional view (rotated 90°), half of the Coat shell is removed to show the 3′ vRNA as well as the rest of the vRNA. **(E)** Secondary structure topology of the Mat^PRR1^, highlighting its two major components: the α-helical region (α-region) and β-sheet region (β-region). Two important β-sheet sub-regions, β4-α1 loop and the tip region, are denoted by arrows. **(F)** The cryo-EM map of the “Mat-less” PRR1 with the Coat shown in pink and the vRNA shown in gray. The lack of Mat density is shown in the inset (viewing angle indicated by the eye cartoon). This class was composed of 67,975 particles (29%) from the PRR1 data-set. **(G)** The Coats (tan) immediately surrounding the Mat^PRR1^ (blue), with the surface contacts between the two labeled red. The surface area of the contacts (803Å^2^) is reported below. **(H)** The Coats (maroon) immediately surrounding the Mat (gray) of MS2 (PDB ID: 5TC1), with the surface contacts between the two labeled red. The surface area of the contacts (1,689Å^2^) is reported below.

In this study, we investigate the interaction between PRR1 and its RP4-encoded T4SS machinery in both *P. aeruginosa* and *E. coli*. We present the mature PRR1 phage structure at 3.45 Å resolution and locate key residues on TrbC that are essential for PRR1 infection. We demonstrate that treatment of the bacteria harboring RP4 plasmids with both infectious and UV-crosslinked, non-infectious PRR1 significantly reduces conjugation. We further found that simultaneous exposure to phage and antibiotic stress selects for RP4 host mutants that fully resist phage infection and dramatically reduce plasmid transfer. A significant subset of mutants—including *traF1, trbH1, trbJ2*, and *trbE1*—exhibited a unique functional “leakiness”, retaining partial conjugation efficiencies (1% to 27% of wild-type levels). These mutants offer a unique opportunity to map the phenotypic landscape of phage sensitivity relative to pilus functionality and be used to reveal the molecular determinants that govern successful ssRNA phage entry.

## RESULTS

### Structure of the Mature PRR1 Virion

Bacteriophage PRR1 is an IncP (RP4)-dependent ssRNA phage. In order to better understand how PRR1 targets IncP pilus and affects host conjugation, we began by determining the structure of the mature PRR1 virion alone using cryo-electron microscopy (cryo-EM) to a resolution of 3.45Å (**Fig. 1D, Fig. S1A-D**). The mature RP4-dependent PRR1 exhibits a pseudo-*T*=3 icosahedral symmetry, with 178 copies of the coat protein (Coat) and a single copy of the Mat forming the capsid.

The capsid of PRR1 has a diameter of 292Å, with the Mat extending 35Å above the surface of the Coat shell. The Mat is composed of two regions angled at 125°: the α-helical region (α-region) and the β-sheet region (β-region) (**Fig. 1E**). The α-region is inserted into the capsid, while the β-region extends outside the virion enabling it to bind to its pilus receptor. Loops connect the two regions, form two layers, with the loop connecting β4 and α1 (termed the β4-α1 loop) being the top layer. The loops in the bottom layer serve as a hinge for the angle between the α-helical and the β-sheet regions. Additionally, the tip region of Mat is composed of loops connecting β-strands to one another within the β-region. The tip region of the Mat^PRR1^ was resolved at poor resolution (**Fig. S1C**), likely due to its high flexibility and is consistent with observations in other ssRNA phages.^17–19^ We also observed the terminal stem loop of the 3′ vRNA extending 20Å outside the capsid, nestling against a groove in the β-region of Mat.

In canonical ssRNA phages MS2 and Qβ, the α-helical regions of the Mat align in such a way that their β-regions face opposite directions.^17,20,21^ Structural comparison shows that Mat^PRR1^ adopts a similar structure and orientation to Mat^MS2^. While Mat^PRR1^ and Mat^Qβ^ have similar α-regions that align with each other, their β-regions orient in opposite directions (**Fig. S1E**). Notably, Mat^PRR1^ displayed a similar structure to Mat^ϕCb5^ except in the tip region, where the two tips curve in opposite directions. Processing of the ∼230,000 PRR1 particles revealed three distinct populations (**Fig. S1A**). Approximately 13% of particles represented mature virions containing both viral RNA (vRNA) and Mat, while the majority (∼57%) consisted of empty capsids, consistent with naturally occurring virus-like particles (VLPs) generated during the assembly process. Notably, a third class, comprising ∼29% of particles, contained defined vRNA density but lacked visible Mat (**Fig. 1F)**. To our knowledge, these “Mat-less” particles with conformationally defined vRNA have not been previously reported for other ssRNA phages. It is possible that PRR1’s Mat has weaker interactions with its surrounding Coats in the capsid, as evident by the smaller area of interface between Mat and the surrounding Coats compared to other phages, such as MS2 (PRR1=803Å^2^, MS2=1689Å^2^ shown in **Fig. 1G** and **H**), potentially leading to its loss during purification. This is also consistent with the previous observation that PRR1 is more RNase sensitive.^22^

### 3′ vRNA Interacts with the Mat

In PRR1, the 3′ terminus of the vRNA folds into a five-stem structure, with the final two stem-loops, U1 and V1, forming a chair-like arrangement that supports the Mat (**Fig. 2A-C**). This conformation constitutes the primary RNA–Mat interaction. We also observed many vRNA-Coat contacts (<5Å RNA/protein distances), facilitating the packaging of the vRNA (**Fig. 2D** and **E**). Notably, within the 3′ end of the vRNA the R1 and U2 loops interact with coat protein dimers (**Fig. 2E** and **Fig. S1F**). These stem loops are “operator-like”, potentially helping to facilitate the packaging of the vRNA through binding of the Coat dimers.^23,24^

**Figure 2.**
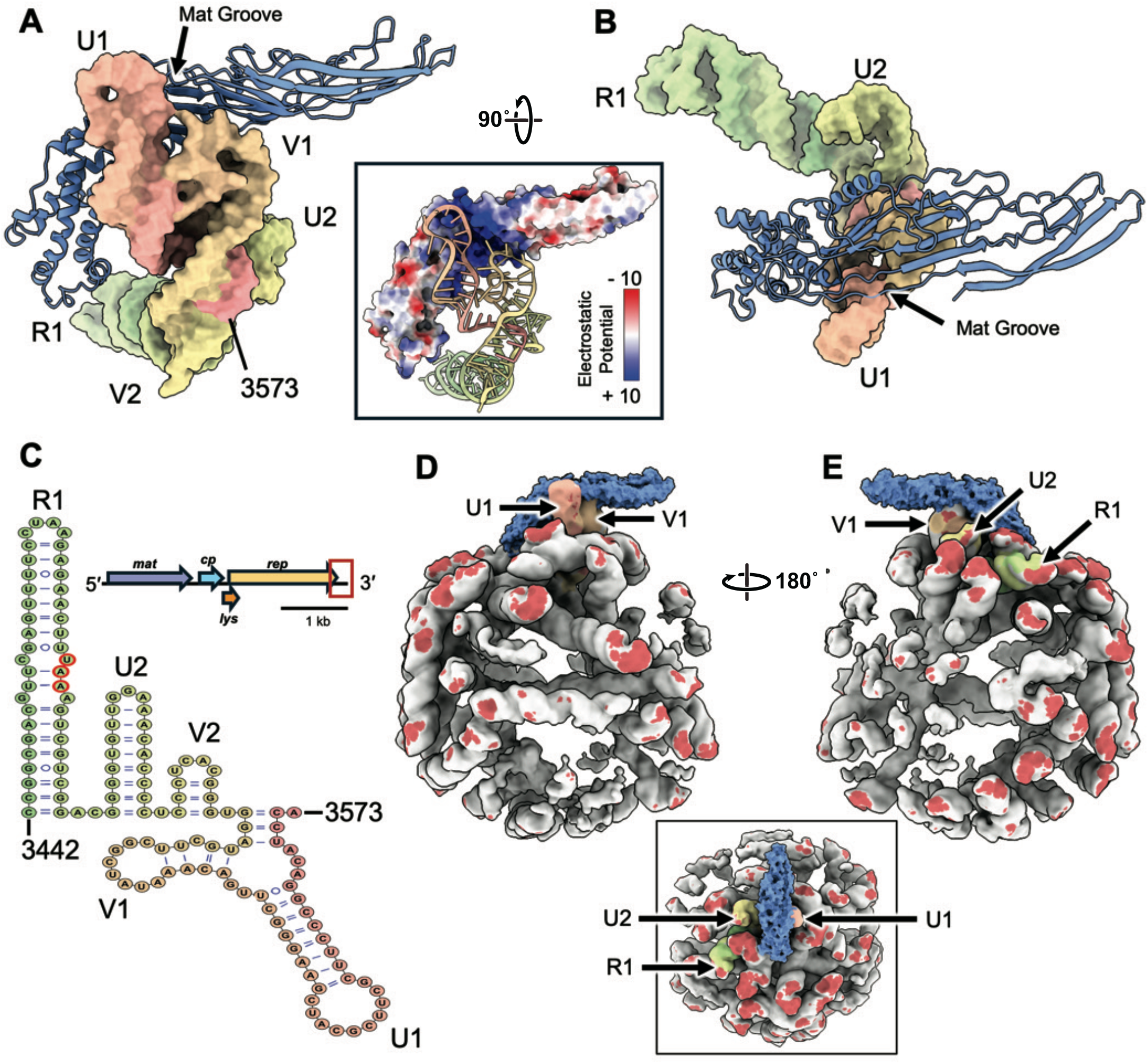
Structural analysis of the vRNA interactions with Mat and Coat. **(A)** The interaction between the Mat and the 3′ end of the vRNA is depicted here, illustrating the vRNA docked into the “groove” of the β-region of Mat. The last nucleotide of the vRNA, A3573, is labeled. The inset shows the electrostatic surface of the Mat^PRR1^, revealing a positively charged surface of the groove that mediates interaction with the negatively charged phosphate backbone of the vRNA. **(B)** The same structure as in Panel **A** but rotated 90°. **(C)** 3′ vRNA (nucleotides 3442–3573) folds into a five-stem-loop secondary structure labeled R1, U2, V2, V1, and U1. The stop codon bases UAA of the replicase gene are indicated by red circles within the R1 stem. The small inset (top right) shows the location of the 3′ vRNA (boxed in red, including R1 and 3′ UTR) in the full PRR1 genome. **(D)** The vRNA and Mat^PRR1^ shown with the Coat shell removed. The 3′ vRNA is colored as in Panels A-C with the rest of the vRNA colored gray. The red touch-ups represent vRNA contacts within 5Å with the Coat shell. The inset shows a top view of the panel, as seen from the Mat. **(E)** The same as Panel **D** but rotated 180° degrees. U1, V1, U2, and R1 helices are labeled in Panels **D** and **E**.

The groove of the β-region in Mat, facing the vRNA, contains a high density of positively charged amino-acid residues, including lysines, arginines, and histidines, which form a strong electrostatic surface (**Fig. 2A**, inset). The negatively charged phosphate backbone of the final RNA stem-loops, U1 and V1, interfaces with this region, stabilizing the Mat within the capsid. Notably, the U1 loop of the 3′ UTR appears partially solvent-exposed while closely apposed to the protein surface (**Fig. 1D**). Similar 3′ UTR architectures have been observed in MS2 and Qβ ^25–27^, and U1–Mat interactions are also conserved in PP7, AP205, and ϕCB5.^18,19,28^ The V1-Mat interaction is novel and may help to stabilize Mat within the virion or to facilitate Mat in escorting the associated RNA into the host cytoplasm during infection.

### Molecular Basis of PRR1 Recognition of the RP4 Pilus

Negative-stain EM confirms the extensive binding of PRR1 to the RP4 pili (**Fig. S2A**). However, due to the low cell-surface abundance—on average, only one pilus is observed per 10 cells— and the short length (200-500 nm) of the RP4 pilus (**Fig. S2B**), we were unable to scale up the purification to obtain sufficient quantities for high-resolution cryo-EM study.

Therefore, we enacted a two-fold approach to pinpoint which residues were involved in the Mat-pilus binding. The T4SS major pilins, of which the RP4-encoded TrbC is a member, are highly conserved. All previously published T4SS pilus structures demonstrate a remarkably consistent C5 symmetry, a rise of 12-15 Å, a twist of 24-30°, and a pilin domain organization (**Fig. 3A**) of α1, α2, loop, and α3_(1/2)_.^29–33^ This conservation ensures that specific regions, namely the N- and C-termini, are always surface-exposed on the filament exterior. As such, generating a structural model of the RP4 pilus was straightforward and reliable, with the measured pitch (15Å) and twist (24°) falling within the reported range for T4SS pili (**Fig. S2C, see Methods**). Furthermore, while phospholipids are a characteristic feature of T4SS pili, their identity varies, and they are consistently localized within the pilus lumen, not facing the exterior, and are therefore unlikely to impact phage adsorption to the pilus surface (**Fig. S2D**).

**Figure 3.**
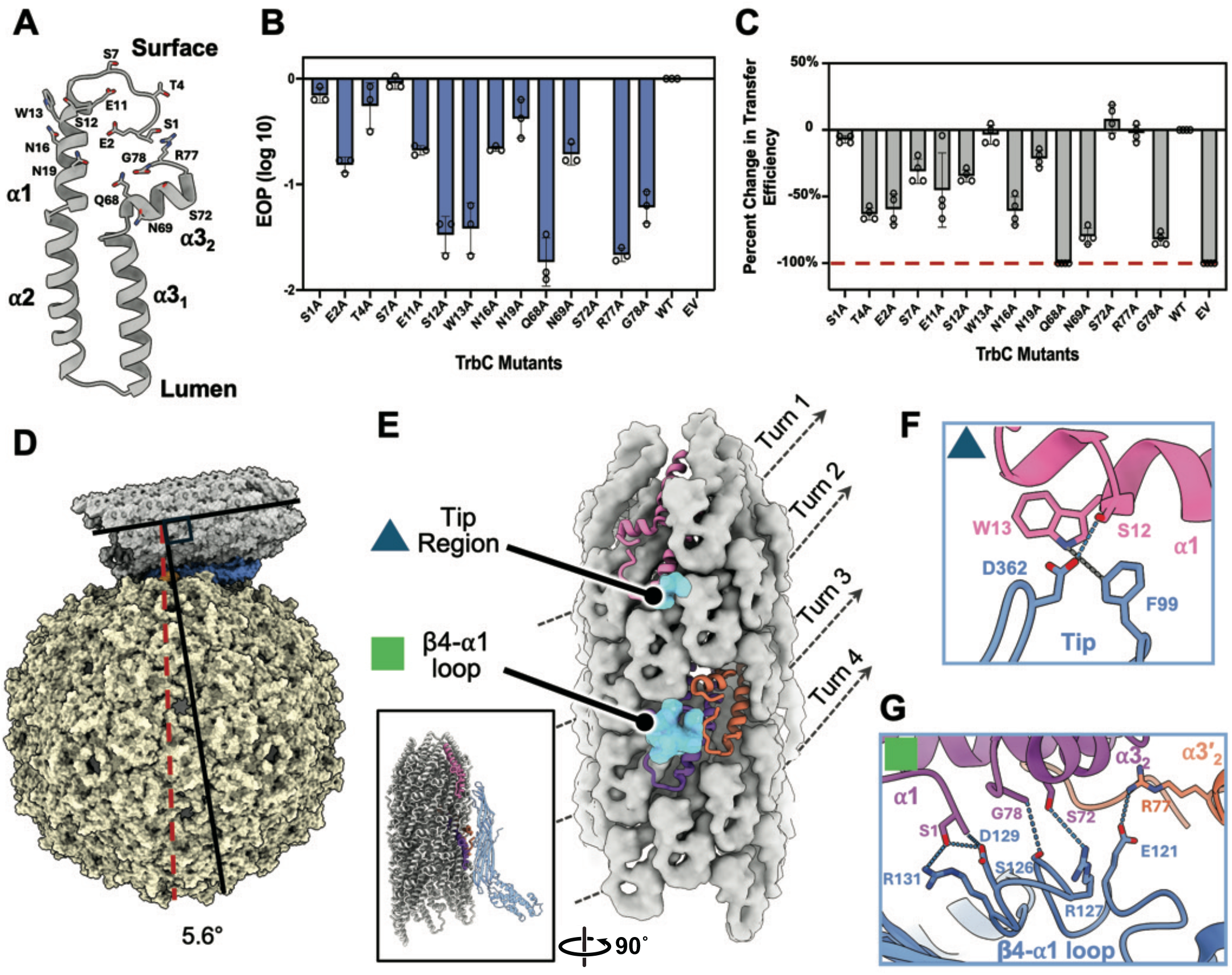
Structural and functional analysis of PRR1 attachment to the RP4 (IncP) pilus. **(A)** Model of a single TrbC pilin monomer with surface exposed residues that possibly interact with Mat^PRR1^ shown in stick representations. **(B)** The change in phage infectivity (titer) of each TrbC mutant is shown as the efficiency of plaquing (EOP). Each bar in the infectivity assay represents the mean ± SD of n=3 biological replicates. The lack of a bar indicates a lack of infection (e.g. S72A). **(C)** The percent change in conjugation efficiencies of TrbC variants carrying mutations in binding residues. Each bar in the conjugation assay represents the mean ± SD of n=4 biological replicates. The dashed red line represents a transfer efficiency of zero. **(D)** The surface representation of the PRR1 virion (tan and blue) bound to the RP4 pilus (gray), using the Mat of the highest-scored docking model of the RP4-Mat complex as an anchor. The axis of the RP4 pilus is shown as a black line with another black line perpendicular to it. A two-fold axis of PRR1 is shown as a red dashed line. The measured tilt angle of binding (between the lines) is 5.6°. **(E)** Side view of the Mat^PRR1^ footprint (blue) on the RP4 pilus (gray). Two Mat regions participate in pilus binding; the β4-α1 loop is bound two TrbC pilin monomers (orchid and orange), and the tip region engages one pilin monomer (light pink). The rest of the pilin monomers (gray) are shown in surface representations. The β-region of Mat^PRR1^ spans three turns on the pilus. The inset shows a 90° rotation of the full Mat bound to the RP4 pilus. **(F)** The tip region (blue shading) from Panel **E** is zoomed in here to show the detailed interactions between the pilus (light pink) and the tip region of the Mat (blue). The gray bond indicates a π-stacking interaction, while the blue line indicates a hydrogen bond. **(G)** The β4-α1 loop (blue shading) from panel **E** is zoomed in here to show the detailed interactions between the pilus (orchid and orange) and the β4-α1 loop of the Mat (blue).

Based on this model of the RP4 pilus, we first performed alanine-scanning mutagenesis of all surface-exposed residues of the major pilin TrbC and assessed mutant susceptibility to PRR1 (**Fig. 3A**). Several mutants of TrbC resulted in a significant drop in PRR1 infectivity, greater than 1 log_10_, including S12A, W13A, Q68A, S72A, R77A, and G78A (**Fig. 3B**). When the pilus function of these mutants was tested, we observed that only mutants S12A, W13A, S72A, and R77A had similar levels of conjugation compared with wild-type, signifying that infectivity loss was not due to a decrease in pilus production but instead due to phage adsorption (**Fig. 3C**). Our mutagenesis was limited to the pilin only, not the Mat^PRR1^, as to avoid potentially compromising genome packaging due to alterations within the RNA sequence.

We then used molecular docking to model the PRR1–RP4 pilus interaction (**see Methods**). We used a density-map–derived model of the Mat^PRR1^ together with our modeled structure of the RP4 pilus. Based on the high structural homology between the MS2-F pili and PRR1-RP4 pili systems at the Mat-pilus interface, we used the Mat^MS2^–F pilus complex (PDB: 6NM5) as a starting reference (**Fig. S2D**).^17^ Similar to MS2, our best scored model (**Fig. S2 E** and **F**) from docking showed that PRR1 bound to the RP4 pilus at a tilted angle (∼5.6°, **Fig. 3D**). Importantly, this model was consistent with the alanine scanning results (**Fig. 3E-G**), with S12_TrbC_, W13_TrbC_, S72_TrbC_, and R77_TrbC_ present at the interface between the RP4 pilus and the Mat. The key interactions localized to two regions of the Mat^PRR1^, the tip region and the β4-α1 loop (**Fig. 3E**). In the tip region, D362_Mat_ from the β9-β10 loop hydrogen-bonded to the sidechain of S12_TrbC_ on the pilus (**Fig. 3F**). We also observed a hydrophobic interaction between F99_Mat_ from the β3-β4 loop and W13_TrbC_, which formed a 3.5Å Π-stacking interaction (**Fig. 3F**). In the β4-α1 loop, E121_Mat_ was seen forming a salt-bridge with R77_TrbC_ (**Fig. 3G**). Our docking also revealed TrbC N- and C-terminal residues S1_TrbC_ and G78_TrbC_ as binding sites for the Mat^PRR1^ (**Fig. 3G**). Although the mutagenesis showed no drop in PRR1 EOP for S1A TrbC mutant, it could be due to the interaction involving the PRR1 Mat and the S1_TrbC_ backbone which was not altered in the alanine scanning mutagenesis; the G78A TrbC mutant was unable to conjugate, likely due to protein misfolding. Specifically in our model of the Mat-pilus complex, the Mat-S1_TrbC_ interactions not only involve sidechain-sidechain hydrogen bonds between S1_TrbC_ and Mat residues R131_Mat_ / D129_Mat_, but also a strong electrostatic interaction between D129_Mat_ with the N-terminal amine of S1_TrbC_ (**Fig. 3G**). Furthermore, R127_Mat_ and S126_Mat_ were both observed hydrogen bonding to the sidechain of S72_TrbC_ and the C-terminal carboxyl group of G78_TrbC_, respectively (**Fig. 3G**).

### PRR1 Abolishes RP4 Mediated Conjugation

Given that the RP4 plasmid drives the spread of antibiotic resistance via conjugation, we investigated whether PRR1 infection could disrupt this process by targeting the RP4-encoded pilus. We performed conjugation assays (**Fig. S3A**) in the presence or absence of PRR1 at various multiplicities of infection (MOIs). In the presence of PRR1, we observed a dramatic reduction in conjugation efficiency, dropping from wild-type levels down to undetectable levels, as we increased MOI (**Fig. 4A** and **Fig. S3B**). Complete inhibition occurred at MOIs ≥1, while only at MOI 0.1 was conjugation retained at a measurable level (>100-fold reduction). We also included ssRNA phage PP7, which is unable to infect this strain of *P. aeruginosa (*Δ*pilA)* due to the deletion of its receptor, the major pilin (PilA) of the type IV pilus, in our conjugation assay as a control for non-specific ssRNA phage interactions. Conjugation with PP7 proceeded at levels similar to our buffer control, which showed that PRR1 specifically interacting with the RP4 pilus is responsible for the reduced conjugation.

**Figure 4.**
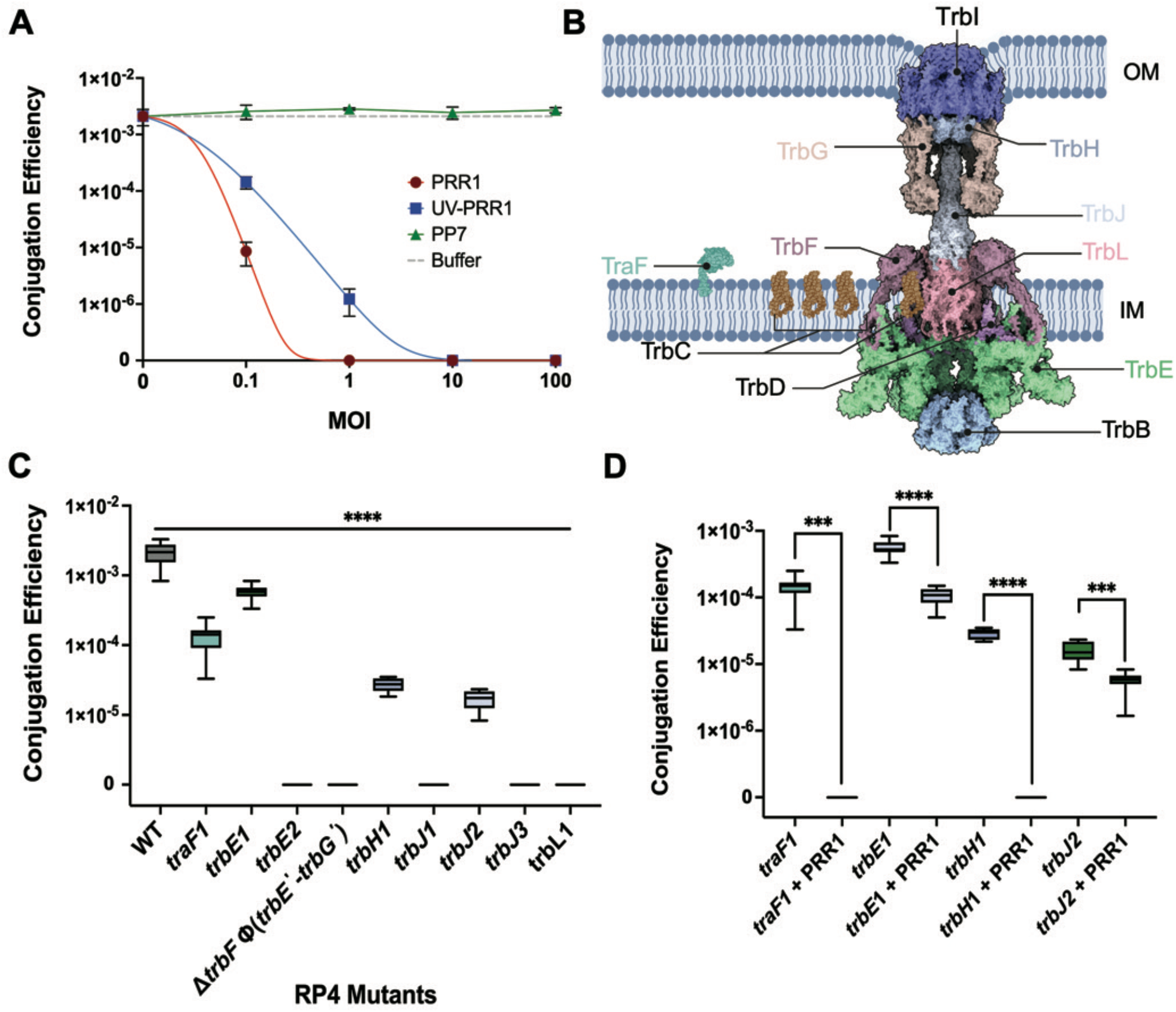
PRR1 reduces conjugation and phage/antibiotic selective pressures force mutations in RP4 core genes. **(A)** The effect of PRR1 (red), UV-PRR1 (blue), PP7 (green), and buffer (gray) on RP4 conjugation in *P. aeruginosa* PAO1Δ*pilA* (RP4). Each point represents the average of n≥4 replicates. **(B)** A model of the RP4 T4SS. The arrangement of the core proteins within the transfer complex is shown. The protein names are colored to indicate that they contain one of the nine mutations. Those left black do not contain a mutation. This structural model of the RP4 system is derived from its homology to the R388 system (PDB: 7OIU, 7O43, 8RT4, 8RT6, 8RT9, 8RTD).^35^ **(C)** The conjugation ability of the nine PRR1 resistant mutants. Data represent mean ± SD of n ≥8 independent replicates. Statistical significance was determined by one-way ANOVA with Dunnett’s multiple comparisons test; all comparisons shown were highly significant (p < 0.0001). **(D)** Conjugation efficiencies of the four transfer-positive mutants in the absence (left) or presence (right) of PRR1 (MOI=10). Data represent mean ± SD of n = 4 biological replicates. Statistical significance was determined using an unpaired two-tailed t-test for each mutant; p-values are: *traF*1= 0.0003, *trbE*1< 0.0001, *trbH*1< 0.0001, *trbJ*2= 0.0002.

We then performed the same conjugation assay in parallel using UV-inactivated PRR1, named UV-PRR1 (**see Methods**), to confirm that the conjugation inhibition was not simply due to host cell lysis or phage replication which depletes cellular resources. UV treatment covalently crosslinks the vRNA to the inner surface of the proteinaceous phage capsid through the contacts we previously observed (**Fig. 2D** and **E**). This renders the phage non-infective (**Fig. S3C**) while preserving its ability to bind pili.^18^ UV-PRR1 treatment produced a similar trend in conjugation inhibition, though at slightly reduced levels at lower MOIs (**Fig. 4A** and **Fig. S3B**). We found that these results hold true in an *Escherichia coli* based RP4 system as well, with PRR1 and UV-PRR1 causing a full reduction in plasmid transfer (**Fig. S3D**). These findings suggest that PRR1 inhibits conjugation primarily through its interaction with the RP4 pilus rather than cell lysis or phage replication.

### PRR1 drives evolution of transfer-deficient RP4 mutants

Beyond characterizing the physical impact of PRR1 on the T4SS, we also investigate the genetic evolution of the RP4 plasmid under phage challenge. The typical bacterial response to a phage targeting a plasmid-encoded receptor (like the RP4 pilus) is the loss of the entire plasmid. To prevent plasmid curing (loss) while promoting specific gene mutations, we implemented a dual-selection strategy: simultaneously challenging bacteria with PRR1 phage and antibiotics, to which the RP4 plasmid confers resistance (**see Methods** and **Fig. S4A**). This approach ensures the retention of the RP4 plasmid due to the essential antibiotic resistance genes yet produces PRR1 resistance (**Fig. S4B**).

The resulting colonies had mutations localized to genes required for T4SS assembly: *traF* in the Tra1 operon and *trbB*-*trbL* in the Tra2 operon (**Fig. 4B**). The T4SS system is defined in five parts: pilins, outer membrane core complex (OMCC), inner membrane core complex (IMCC), stalk, and ATPases. The major pilin subunit is known to be TrbC. TrbJ is the minor pilin subunit, which initiates the assembly of TrbC filament.^14,34,35^ The TraF peptidase is needed to mature TrbC, cleaving on the C-terminal side of TrbC.^16^ The OMCC is predicted to be comprised of TrbGHI, forming a pore structure.^35,36^ The IMCC is composed of TrbD, TrbL, a single helix of TrbE, and the tail portion of TrbF, which is imbedded in the membrane.^36,37^ The stalk spans the periplasmic region and is composed of TrbF, TrbJ, and TrbL, with TrbL forming the core of the complex and containing the sites of major pilin assembly.^34–36^ The ATPases are necessary for pilus extension and retraction, and this system is known to contain two of them: TrbE and TrbB.^37,38^ TrbE is known to form a dodecamer that has a single transmembrane helix inserted in the inner membrane, connecting the IMCC to the cytosol where the hexameric TrbB resides. TrbB and TrbE are thought to interact to coordinate pilus extension and retraction.^35,38,39^

In total, we have identified nine mutations that confer PRR1 resistance and exclusively in the RP4 T4SS machinery. No mutations were found on the bacterial genome, with most being in-frame insertions or deletions in core T4SS genes; one mutant contained a frameshift in *trbE*. These genes are responsible for biogenesis and dynamics of the RP4 pili^12,36^, which are essential for a successful PRR1 infection. To confirm that resistance was plasmid-encoded, each mutant plasmid was transformed into a native wild-type PAO1 host before being challenged with PRR1. In all cases, PRR1 resistance was retained, confirming that the phenotype was plasmid-borne (**Fig. S4C**).

### Transfer Efficiency Varies Among PRR1-Resistant Mutants

We next assessed the conjugation efficiency of each mutant. Five mutants (*trbE2*, Δ*trbFϕ(trbE′-trbG′)*, t*rbJ1, trbJ3*, and *trbL1*) completely lost the ability to transfer (**Fig. 4C** and **Fig. S4D**), which is consistent with previous results that mutations within the core T4SS machinery abolish transfer.^12^

Strikingly, four remaining mutants retained some conjugation ability despite full resistance to PRR1. Three of these mutants, *traF1* (ΔA19-G25), *trbH1* (ΔQ76-A108), and *trbJ2* (R234_Q235 ins QAQQDR) (**Fig. S5A-C**), were able to transfer at extremely low rates: all below 7% of the wild-type conjugation rate (**Table S1, Fig. 4C**, and **Fig. S4D**). However, the last mutant, *trbE1*, had reduced transfer, but at a significantly higher level than the previous three: ∼27% of the wild-type conjugation efficiency (**Table S1, Fig. 4C**, and **Fig. S4D**). Conjugation assays using native wild-type PAO1 hosts, with mutant plasmids transformed, showed consistent conjugation efficiencies compared to the RP4 mutants, which we originally screened from our phage resistance assays (**Fig. S4 E** and **F**).

Interestingly, *trbE1* contained a 10 base pair deletion resulting in a frameshift at amino acid 134. This caused 18 non-wild-type amino acids to be translated, ending in a premature stop codon at position 151 (**Fig. S5D**). When further analyzing the coding sequence of *trbE1*, we found a downstream alternative start codon that may produce a shorter isoform; this N-terminus-removed truncation spans amino acid 146 to the wild-type stop codon. Additionally, a multiple sequence alignment of TrbE homologs revealed this conserved internal start codon (**Fig. S5E**). We found literature supporting this observation, as a previous study reported co-purification of a ∼70 kDa TrbE isoform along with the full-length TrbE (94 kDa)^37^; this matches the predicted mass of the protein product from the secondary start site. This shorter product was shown to be conjugation deficient.^37^ Therefore, it is possible that our mutant TrbE needs both protein products, Residues 1-151 and Residues 146-852, to mimic the wild-type TrbE protein structure and function, potentially through the formation of a heterodimer (**Fig. S5F**).

### PRR1 Further Inhibits Conjugation in Resistant Mutants with Reduced Transfer Efficiency

Surprisingly, all sequenced mutants generated via our laboratory evolution that exhibited phage/antibiotic resistance had mutations located outside the *trbC* gene, which encodes the major pilin protein that directly interacts with the phage during adsorption. Particularly, for the four phage-resistant mutants that retain partial conjugation, the wild-type TrbC pilin protein is presumably expressed to form RP4 pili to interact with PRR1.

We then test whether PRR1 could further suppress conjugation in these PRR1-resistant mutants that retained partial transfer activity. Conjugation assays in the presence of PRR1, MOI of 10, revealed that *traF1* and *trbH1* mutants showed a complete loss of transconjugants, while *trbE1* (94.9%) and *trbJ2* (99.7%) exhibited significant additional reductions in transfer (**Fig. 4D, Table S1**, and **Fig. S4G**). Despite being resistant to phage infection (**Fig. S4B**) these mutants appear to still produce pili capable of interacting with PRR1. Therefore, the adsorption of PRR1 and its potential interactions with the T4SS assembly machine is sufficient to inhibit conjugation.

## DISCUSSION

In this study, we present the structural and functional characterization of the ssRNA phage PRR1 and its interactions with the conjugative IncP plasmid RP4. Using cryo-EM, we resolved the mature PRR1 virion at 3.45 Å resolution and identified key structural features that illuminate conserved and divergent vRNA packaging strategies among ssRNA phages. Specifically, Mat^PRR1^ shows a strikingly similar structure to Mat^MS2^, with a conserved U1-Mat interaction. However, the V1-Mat interaction is novel and not present in other ssRNA phages we studied so far. It most likely contributes to stabilizing the Mat within the capsid or facilitating the delivery of the attached vRNA.

Unexpectedly, we resolved a subpopulation of particles lacking the Mat, suggesting dissociation from the virion. This observation, together with the partially solvent-exposed U1 loop of the 3′ UTR, could explain historical PRR1 RNase sensitivity.^22^ The absence of Mat in this class may result from weaker interactions between the Mat and Coat proteins potentially allowing for dissociation of the Mat post-assembly.

Our alanine screening on TrbC major pilin identified key residues mediating PRR1 attachment to the RP4 pilus, revealing an interface largely driven by hydrogen bonding, electrostatic interactions, and a hydrophobic interaction (F99_Mat_ and W13_TrbC_ Π-stack). Importantly, some of these mutants maintained conjugation efficiencies comparable to wild-type RP4, indicating that the drop in infectivity observed is not due to lack of pilus production or function, but rather due to compromising the phage binding interface. The modeling of the binding interaction supported the results of the alanine mutagenesis and identified N- and C-terminal residues S1_TrbC_ and G78_TrbC_ as additional binding residues via backbone amine and carboxyl groups respectively.

PRR1 significantly inhibited RP4-mediated conjugation, even when UV-inactivated and thus incapable of infection. In our previous work, we discovered that UV-inactivation of PP7, another ssRNA phage that infects *P. aeruginosa* via type IV pili, resulted in the same level of pilus detachment as wild-type PP7.^18^ Through this we know that it is binding and subsequent pilus retraction alone that results in pilus detachment. Therefore, this along with the fact that MS2 detaches conjugative pili upon infection leads us to hypothesize that UV-PRR1 is acting in a similar manner to cause reduction in conjugation due to a direct impact on the T4SS, potentially through pilus removal or jamming of the T4SS. Alternatively, an overload of phage particles on the T4SS may create too large of a force barrier to overcome causing the RP4 pilus to lose its ability to retract, thus blocking conjugation. Both hypotheses are yet to be tested.

The significance of this work comes in the form of PRR1’s dependence on RP4 (IncP), which has a much broader host range than other incompatibility plasmids. Therefore, the use of PRR1 to inhibit the spread of plasmid-based antibiotic resistance is relevant as it’s more applicable to a wider range of pathogenically relevant bacteria.^22^ These findings also raise the possibility of developing Mat-derived peptide inhibitors, a concept supported by recent work showing that isolated Mat domains can inhibit pili.^28,40^

Our experimental approach—simultaneously challenging bacteria with PRR1 phage and antibiotics (to which the RP4 plasmid confers resistance)—was highly effective in forcing the evolution of specific phage-resistance mutations while ensuring the retention of the target plasmid (RP4). This strategy provided a direct, first-hand model of the evolutionary pressures exerted in a phage-antibiotic synergy (PAS) scenario.^41^ In many PAS studies, the combination of phage and antibiotic is used to select for phage-resistant mutants that, as a pleiotropic effect, become hypersensitive to the antibiotic. This often occurs because the mutation conferring phage resistance simultaneously compromises a critical bacterial function, such as maintaining membrane integrity or effective efflux^42,43^, thereby reducing fitness or enhancing antibiotic uptake. Our finding that the dual-selection strategy exclusively drove mutations to T4SS-associated genes on the RP4 plasmid, and not to other genes in the bacterial chromosome, demonstrates the power of this method to isolate highly targeted genetic changes. Furthermore, the discovery of nine such mutants provide a valuable toolkit for future PAS investigations. Specifically, the fact that eight of the nine mutants lost conjugation or had severely impaired conjugation capability is significant. While these mutants may lose the ability to spread AMR via HGT, it also suggests that the mutation that caused phage resistance came at a high fitness cost to the plasmid function.

However, the one remaining TrbE mutant, which maintained ∼30% conjugation efficiency, is particularly compelling. This partial function is hypothesized to result from a functional TrbE complex formed by two separate truncated products—one N-terminal (ends at a premature stop codon due to a frameshift mutation) and one C-terminal (produced from an alternative start site). This may represent a novel phage resistance mechanism achieved by remodeling the assembly and stability of the T4SS while maintaining partial conjugation ability. The survival of this T4SS-remodeling mutation with reduced fitness cost suggests that the selective pressure of PAS can drive bacteria to find “smart” resistance mechanisms that minimize functional impairment. Studying this partially functional T4SS complex provides critical insight into the T4SS’s modularity and stability and how bacteria might evolve phage resistance without sacrificing the key fitness advantage of conjugation (and thus, AMR spread) in a natural environment.

## METHODS

### Bacteriophage and Bacterial strains

All *E. coli* and *P. aeruginosa* strains were cultured in LB medium (BD Biosciences) containing 10 g/L Bacto tryptone, 5 g/L Bacto yeast extract, and 10 g/L NaCl. Bacterial strains were initially streaked from permanent glycerol stocks onto LB agar (LB supplemented with 15 g/L Bacto agar) and incubated overnight (16 h) at 37 °C. A single colony from freshly-streak plate was then inoculated into 5 mL of LB medium and grown overnight at 200 rpm. When required, LB broth or LB agar was supplemented with 50 µg/mL kanamycin (LB+Kan), 100 µg/mL tetracycline (LB+Tet), 60 µg/mL gentamicin (LB+Gm), 30 µg/mL chloramphenicol (LB+Cm), or 300 µg/mL carbenicillin (LB+Cb). The bacteriophages and bacterial strains used were summarized in **Table S2**.

### PRR1 Production

PRR1 was originally produced from a pRSF_PRR1 plasmid. The PRR1 genomic DNA was synthesized based on the DNA sequence from NC_008294 by Genscript. The synthesized PRR1 DNA genome was cloned into pRSF plasmid. The plasmid was transformed into BL21(DE3)pLysS for phage production with IPTG induction at a final concentration of 2mM overnight at 37°C. The supernatant from BL21(DE3)pLysS was titered onto *P. aeruginosa* PAO1Δ*pilA* (RP4); a strain lacking type IV pili making it resistant to ssRNA phage PP7 and containing the RP4 plasmid for T4SS production. A single PRR1 plaque was picked and mixed with 50 µl of overnight PAO1Δ*pilA* (RP4) in 5 ml LB+Tet. The infection was allowed to go overnight, shaking at ∼200 rpm at 37℃. This entire 5 ml lysate was then used to infect a 200 ml culture of the host at an OD= 0.6. The infection was allowed to go overnight, shaking at ∼200 rpm at 37℃. The entire 200 ml lysate was used to infect an OD=0.6 3L culture of the host, which was allowed to go overnight, shaking at ∼200 rpm at 37℃. The overnight culture was spun at 6,000 ×g for 40 minutes to pellet cells. The supernatant was then filtered with bottle top vacuum (0.45µm PES) filter 0.45µm (Corning). After each expansion step a spot titer was performed. The resulting titer was approximately 10^10^ -10^12^ pfu/ml. The filtered supernatant was subjected to tangential flow filtration using a VivaFlow 200 (Sartorius) cross-flow filtration with a 100 kDa molecular weight cut off to concentrate the supernatant. The supernatant was concentrated to ∼40 ml. Since it was concentrated, the solution contained a high concentration of *P. aeruginosa* endotoxin that needed to be removed. The following protocol was followed.^44^ Briefly, an equal amount of supernatant and octanol were mixed together for 10 minutes, centrifuged at 4000 rpm to induce phase separation, and the top aqueous phase was saved. This was repeated several times until the interface layer was minimal. The titer after endotoxin removal was usually 10^10^ -10^12^ pfu/ml. The remaining solution was then clarified by adding 1 unit/ml of DNAse I (NEB), incubating at room temperature for 1 h, and centrifuging at 10,000 ×g for 1 h at 4°C. The clarified solution was then used for CsCl isopycnic density ultracentrifugation. A CsCl was gradually added to the phage solution to achieve the density of 1.4 g/ml and the solution was spun at 45,000 rpm with a Ti40 rotor at 4°C for 24 h. The resulting bands were collected and dialyzed against 1 L high-salt buffer (50 mM Tris pH 8.0, 1 M NaCl buffer) in a 20 kDa MWCO Slide-A-Lyzer™ G2 Dialysis Cassette overnight, followed by 4 L of phage buffer (50 mM Tris pH 8.0, 150 mM NaCl) overnight two times at 4℃. The resulting titer of phage after this final step was regularly 10^11^ -10^13^ pfu/ml.

### Cryo-EM Sample Preparation and Data Acquisition

Cryo-EM grids of PRR1 were prepared by applying 5 µl of the purified phage to C-flat R2/1 300 mesh Cu grids that were glow-discharged for 60 sec with a PELCO easiGlow™. The grids were blotted and plunge-frozen using EM GP2 Automatic Plunge Freezer (Leica Microsystems). Data acquisition was performed on a Titan Krios G4i microscope (Thermo Fisher Scientific) equipped with a Gatan BioContinuum energy filter (silt width, 15 eV) at 300 kV. Data was collected on a K3 Summit direct detection camera (Gatan) in super resolution mode, yielding a pixel size of 0.86Å after binning by 2. The total of 4,397 movies were collected with 2.5s exposure over 40 frames, resulting in a cumulative dose of 50 e^-^/Å^2^. The data collection parameters were summarized in **Table S3**.

### Cryo-EM Data Processing

The total of 4,397 movies were processed using cryoSPARC v4.5.1. The movies were subjected to patch motion correction and patch CTF estimation. Only the micrographs with CTF-fit resolution better than 7.5 Å were selected, yielding 4,321 micrographs. The PRR1 particles were picked by template-picker with 2D templates generated from a subset of particles picked from blob picker. A total of 232,842 particles remained after selection from good 2D class averages. The particles were subjected to ab-initio reconstruction with 4 classes, yielding a good class of capsid with vRNA density containing 99,167 particles. After non-uniform refinements and 3D classifications, 67,975 particles were classified as “Mat-less” and 30,442 particles were classified as mature. The mature particles were further subjected to 3D variability analysis to glean more Mat density. The data processing pipeline was summarized in **Fig. S1A**.

### Model Building and Refinement

In order to model the PRR1 capsid, we used the previously solved crystal structure of the PRR1 virus-like particles (VLP, PDB code 2VF9).^45^ The structure was refined into the coat density using Phenix real-space refinement.^46^ To model the Mat^PRR1^, the Mat density was segmented out from the refined map using Segger.^47^ We used AlphaFold to predict the model of the Mat^PRR1^ from its sequence (NC_008294). The α-region of the Mat^PRR1^ was first fit and refined into the map using Coot^48^, ISOLDE^49^, and Phenix real-space refinement. Due to the poor-resolution at the β-region, the tip was flexible-fitted using ISOLDE^50^ into the density map that was gaussian low-passed to standard-deviation of 1.2 using ChimeraX.^51^ The same refinement process was repeated for the full Mat.

To model the 3′ UTR and R1 of the vRNA we used the density map corresponding to the mature virion. The strategy used here is similar to the ones we used for Qβ and PP7.^18,25^ We first used windows of about 100 nucleotides to predict the minimum-free-energy secondary structure of the 3′ vRNA.^52^ Then, we modeled RNA structure into its density using Rosetta RNA Denovo^53,54^. The model was further refined with Rosetta RNA ERRASER^55^ and Phenix refinement. Due to the heterogeneity and weak vRNA density before the 3′ region of the vRNA, we were unable to map the rest of the vRNA sequence into our density.

### Alanine Scanning Mutagenesis

First, the wild-type *trbC* gene was cloned into the pUCP19 plasmid backbone for overexpression in *Pseudomonas aeruginosa* strain PAO1Δ*pilA* (RP4Δ*trbC*). To introduce each alanine point mutation into the *trbC* gene, DNA of 150 basepairs was synthesized for each construct (IDT). Inverse PCR was performed using the pUCP19_*trbC* plasmid, and the short inserts were ligated into this vector using HiFi assembly (NEB). The resulting plasmids were transformed, extracted, and sequenced to reveal correctly assembled plasmids. The plasmids were then transformed into the PAO1Δ*pilA* (RP4Δ*trbC*) strain. We then used these strains to test phage infectivity through a titer and pilus function through a conjugation assay. Calculations for EOP of each TrbC mutant were defined as: log_10_(pfu/ml on mutant/pfu/ml on wild-type). Calculations for the percent change in transfer efficiency were defined as: (transfer efficiency of mutant/transfer efficiency of wild-type)×100. The transfer efficiency is defined below in the **Conjugation Assay** section.

### Mat-Pilus Docking

To model the interaction between the Mat of PRR1 and the RP4 pilus, a model of the pilus first needed to be generated. AlphaFold was used to construct a 50-mer structure of the pilus. We then used our model of the Mat^PRR1^ solved from our cryo-EM density map. Since the Mat proteins of PRR1 and MS2 were structurally similar (**Fig. S1E**), the MS2-F-pilus complex was used as a starting point for Mat docking. Match-maker was used to align the two Mats and individual subunits of the F and RP4 pili. Rosetta Dock local docking^56^ was used to perform the docking and 10,000 structures were generated. The score and RMSD for each structure were graphed (**Fig. S2E**) and the ten best structures were visualized in ChimeraX (**Fig. S2F**).^57^ The top structure was further analyzed based on its contacts, hydrogen bonds, salt bridges, and hydrophobic interactions.

### UV treatment of PRR1

The previously purified PRR1 was diluted to a concentration of 10^10^ pfu/ml and aliquoted into 200 μl thin-walled PCR tubes. These samples were then placed directly underneath a germicidal UV lamp for one hour. The titer of the phage was determined via conventional spot titer assay.

### Conjugation Assay

This assay was adapted from Gordils-Valentin *et. al*.^13^ An overnight growth of donor, PAO1Δ*pilA* (RP4), and recipient, PAO1Δ*pilA* (pUCP19*_*GmR), or in the *E. coli* assay donor, XL1-Blue (RP4) and recipient, XL1-Blue (pUCP19_CmR), were used to inoculate 5 ml overday cultures with appropriate antibiotics (*P. aeruginosa*-donor= tetracycline 100 µg/ml, recipient= 60 µg/ml gentamicin) (*E. coli*-donor= kanamycin 50 µg/ml and recipient= chloramphenicol 25 µg/ml). Each culture was grown to an OD_600nm_ of 0.5 and then spun at 4000 rpm to pellet the cells. The cells were washed three times with LB broth to remove any antibiotics. Cells were then pelleted at 10,000 ×g for 5 minutes. Afterwards, the donor and recipient cells were mixed in a 3:1 ratio. If phage was added to the assay, it occurred at this step. 50 µl of the cell mixture was spotted on a Whatman™ 0.45µm cellulose acetate membrane and placed onto a LB agar plate. The plate was incubated at 37℃ for 2.5 h and the membrane was removed from the plate. To remove the cells from the membrane, it was placed in a 50 ml falcon tube with 1 ml of LB broth and vortexed at max speed for 2 minutes. The resulting solution was then spun at 17,000 ×g for 5 minutes to pellet the cells. They were resuspended in 50 µl LB and a 10-fold serial dilution was performed. 5 µl of each resulting dilution was then spotted on LB agar plates containing 100 µg/ml of tetracycline and 60 µl/ml of gentamicin (for *P. aeruginosa*), or kanamycin 50 µg/ml and 25 µg/ml chloramphenicol (for *E. coli*). The plates were then incubated at 37℃ for 16 h. The colony forming units (CFUs) were then counted using the following formula: (# of colonies counted ×10^# of dilutions^/5µl)×50 µl. The conjugation efficiency was calculated from this using the following formula: CFU of transconjugants/CFU of total recipients. The percent reduction as shown in **Table S1** was calculated using the following formula: (conjugation efficiency of mutant/conjugation efficiency of wild-type)×100.

### Isolation of PRR1 Escape Mutants

The RP4 mutants were created through a full plate titer of PRR1 with PAO1Δ*pilA*(RP4), on LB + 100 µg/ml tetracycline, 300 µg/ml carbenicillin, and 100 µg/ml kanamycin. The plate was incubated at 37°C for 48 h to allow PRR1 resistant mutants to spontaneously grow. Colonies that grew within the plaque clearings from PRR1 were picked and streaked on a LB agar plate containing 100 µg/ml of tetracycline. Individual colonies were then selected from each plate and inoculated in 5 mL of LB media and incubated overnight at 37°C with shaking at 190 rpm. The resulting growth was used for total DNA isolation using a Monarch® Spin gDNA extraction kit. A small amount of the growth was used to confirm that the colonies were resistant to PRR1 using a spot titer assay. The RP4 plasmid and bacterial genome extracted were sequenced by Plasmidsaurus. This was done for ten colonies in total. The resulting fasta files were aligned with either the reference plasmid (RP4) or PAO1 genome using NCBI BLAST^58,59^ and Geneious.^60^

To confirm that the RP4 mutant plasmids were the source of the resistance to PRR1 and not a mutation in the host’s genome, the extracted mutant plasmids were transformed into a native wild-type PAO1 host strain: PAO1. The protocol developed by Choi, Kumar, and Schweizer was used to make *P. aeruginosa* electrocompetent.^61^ Approximately 500 ng of mutant RP4 DNA was used to electroporate PAO1. The transformed cells were allowed to recover at 37°C for 2 h, and then they were streaked on a LB agar plate containing 100 μg/ml of tetracycline. Several colonies from each mutant were picked and grown. These mutant colonies were then subjected to a spot titer with PRR1 to confirm resistance and a conjugation assay to compare their efficiency to the first round of mutants.

### Statistical analysis

Statistical analysis of the conjugation assay data was performed using GraphPad Prism’s built in analysis tool (RRID: SCR_002798). A one-way ANOVA was used to determine significance for the mutant comparison followed by a Dunnet’s test for multiple comparisons (**Fig. 4C**). To determine the significance of the conjugation assays with native PAO1 hosts, a one-way ANOVA was used followed by Sidak’s test for multiple comparisons (**Fig. S4F**). For the statistical analysis of the transfer positive mutants with and without PRR1 addition (**Fig. 4D**), an unpaired two-tailed Student’s T-test was used.

## ACKNOWLEDGEMENTS

We are grateful to Ryland Young at the Center for Phage Technology for his gift of the RP4 plasmid. We thank Dr. Gaya Yadav from the Texas A&M University Laboratory for Biomolecular Structure and Dynamics (LBSD) for assistance in cryo-EM data collection. The department of Biochemistry and Biophysics, AgriLife, Texas A&M University, and the Cancer Prevention and Research Institute of Texas (CPRIT) jointly support the LBSD. We also thank the Microscopy and Imaging Center at Texas A&M University, for their support in negative stain-EM. Furthermore, we would like to thank the Texas A&M High Performance Research Computing Center for providing the computational resources required for data processing. We thank Lianna Lill for commenting on the earlier version of this manuscript.

## Funding

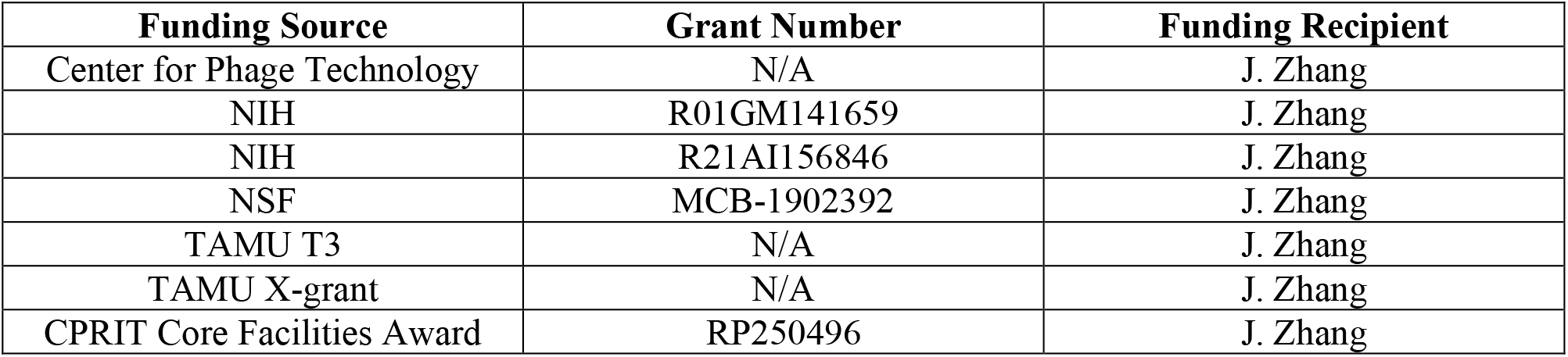

## Author Contributions

Conceptualization: Z.L, J.T., and J.Z. Investigation: Z.L., J.T., D.S., and J.Z. Funding acquisition: J.Z. Supervision: J.Z. Writing – original draft: Z.L., J.T., and J.Z.. Writing – review and editing: Z.L., J.T., D.S., and J.Z.

## Data and Materials Availability

The coordinates and cryo-EM map for the PRR1 virion are deposited in the Protein Data Bank (PDB) and the Electron Microscopy Data Bank (EMDB) with the accession codes: 9ZZY and EMD-75020, respectively.

## Supplemental Materials

Supplemental data: Figures S1-S5 and Tables 1-3

